# Loss of Copine D Leads to Ras Activation in *Dictyostelium discoideum*

**DOI:** 10.64898/2026.04.08.713907

**Authors:** Cody T. Morrison, Sela K. Damer-Daigle, Allison G. Maillette, Cynthia K. Damer

**Affiliations:** Biochemistry, Cell & Molecular Biology Program, Central Michigan University, Mount Pleasant, Michigan, United States of America; (C.T.M.); (C.K.D.); Department of Biology, Central Michigan University, Mount Pleasant, Michigan, United States of America; (S.K.D.D.); (A.G.M.)

## Abstract

Copines are a family of calcium-dependent phospholipid-binding proteins conserved across numerous eukaryotes. The expression of multiple copine genes is dysregulated in several human cancers, yet their molecular functions remain poorly understood. We are investigating copine genes (*cpnA-cpnF*) in the amoeba *Dictyostelium discoideum*, a well-established model for studying conserved signaling pathways that regulate chemotaxis. During development, individual amoebae release and move toward cAMP. cAMP chemotaxis utilizes Ras/PI3K signaling, a pathway that is highly conserved in mammalian cells and often dysregulated in cancer. Using two *cpnD* mutants that were generated via restriction enzyme-mediated integration (REMI), we found that *cpnD* mutants exhibited increased cellular proliferation, precocious development, and larger fruiting bodies. Additionally, we found that *cpnD* mutants had increased cell spreading, producing a flattened morphology and larger cell size. Because activated Ras has been shown to promote this phenotype, we measured Ras activity and found that *cpnD* mutants exhibited increased Ras activation. *cpnD* mutants also formed significantly smaller contractile vacuoles during osmotic stress. Inhibition of PI3K suppressed both the enlarged cell size and reduced contractile vacuole phenotypes, indicating that these defects result from increased Ras/PI3K signaling. Finally, we found that GFP-tagged CpnD transiently localized to the plasma membrane following cAMP stimulation, suggesting CpnD may have a role in cAMP signaling. Together, these findings identify CpnD as a potential negative regulator of Ras signaling and provide the first evidence of copine involvement in the Ras/PI3K pathway, suggesting a conserved mechanism that may help explain how altered copine expression contributes to cancer progression.

## 1. Introduction

Copines are a family of evolutionarily conserved, calcium-dependent membrane-binding proteins found in a variety of eukaryotes ranging from unicellular organisms to humans [1]. Copines are characterized by a conserved domain structure consisting of two N-terminal C2 domains and a C-terminal A domain that is similar to the von Willebrand factor A (VWA) domain found in integrins [1]. The C2 domains function to facilitate calcium-dependent lipid binding [1,2], whereas the A domain serves as a protein-binding motif [2]. Together, these domains suggest that copines act as calcium-responsive sensors that recruit or regulate proteins at membrane surfaces in response to intracellular calcium signals [2]. Humans have nine copines encoded in their genome that exhibit differential expression throughout various tissue types. Dysregulated copine expression has been reported in numerous cancers and human diseases [3–8]. Additionally, functional studies suggest that human copines play a role in promoting cell proliferation, migration, and metastasis [8–10]. Despite these associations, the molecular and cellular mechanisms by which copines regulate signaling pathways remain incompletely understood, highlighting the need for further investigations of copine function in vivo.

To address this, our lab aims to define the mechanistic function of copine proteins using the model organism *Dictyostelium discoideum*. *Dictyostelium* has six copine genes, while other model organisms either have no copine genes, like yeast and *Drosophila*, or only a few copine genes, like *C. elegans* [11]. In their vegetative state, *Dictyostelium* exists as single-celled amoebae that feed on bacteria through phagocytosis; however, upon nutrient depletion, changes in gene expression initiate *Dictyostelium* to undergo development. Development begins when starving amoebae release cyclic adenosine monophosphate (cAMP) pulses, which act as a chemoattractant to initiate aggregation. Cells aggregate into mounds, which transform into migrating slugs. The slugs culminate into fruiting bodies that consist of a stalk supporting a spore head [12]. Early developmental aggregation involves changes in cell migration and adhesion, both of which are processes important in cancer cell metastasis. Therefore, by leveraging *Dictyostelium’s* unique biology, we aim to uncover the conserved copine functions relevant to both normal cellular processes and cancer progression.

*Dictyostelium* aggregation is a well-studied model of chemotaxis, where individual starving amoebae detect, relay, and move toward periodic pulses of cAMP. Chemotaxis is driven by a complex signal transduction network that converts the external cAMP signal into localized actin polymerization at the leading edge of the cell. Extracellular cAMP binds to G-protein-coupled receptors (cAR1), which activate heterotrimeric G-proteins *Ga2bg*. This leads to the rapid activation of Ras proteins at the leading edge of migrating cells [13]. Ras signals to downstream effectors like phosphoinositide 3-kinase (PI3K) to drive signaling cascades that are involved in the regulation of cell polarity, chemotaxis, F-actin dynamics, phagocytosis, and gene expression [14–17]. Studies have shown that constitutively active RasC or RasG leads to hyperactivation of signaling pathways, resulting in increased cell spreading, cytoskeletal activity, and altered migration [18–22]. Importantly, the mechanisms of Ras-mediated chemotaxis are conserved between *Dictyostelium* and mammalian cells. In mammalian cells, Ras signaling pathways also regulate cellular proliferation, growth, survival, and metabolism, and are frequently activated by mutations in cancer [23,24]. As such, the Ras signaling pathway is often targeted for therapeutic interventions in cancer models [25].

The six copine genes (*cpnA-cpnF*) that encode proteins CpnA-CpnF in *Dictyostelium* exhibit 28-60% amino acid identity and have distinct expression patterns throughout development, suggesting that they perform nonredundant functions [11,26]. Studies with *cpnA* null (*cpnA-*) cells have revealed numerous cellular phenotypes, including defects in chemotaxis, adhesion, cytokinesis, contractile vacuole (CV) function, development, and phosphatidylserine (PS) exposure [26–30]. *cpnA-* cells also showed increased cytosolic calcium concentrations, indicating a role for CpnA in calcium homeostasis [30]. *cpnC-* cells exhibited different cellular phenotypes, including distinct defects in chemotaxis, adhesion, and development. *cpnC-* cells also exhibited significantly reduced expression of RegA, a cAMP phosphodiesterase, important in the regulation of intracellular cAMP levels [31]. In this study, we describe the phenotypic analysis of *cpnD* mutants. Here, we show that *cpnD* mutants display distinct defects in morphology, growth, development, and CV size, along with increased Ras activation. Furthermore, we show that GFP-tagged CpnD localizes to the cytosol and transiently translocates to and from the plasma membrane in response to cAMP stimulation. Together, all three copine proteins appear to serve as regulators of key cell signaling pathways.

## 2. Materials and Methods

### 2.1. Dictyostelium cell strains and culture

The parental axenic strain (AX4) and the *cpnD* mutants were obtained from the *Dictyostelium* Stock Center [32]. AX4 and *cpnD* mutants were grown on plastic Petri dishes at 18 °C in VL-6 media (Formedium, VL60102) supplemented with penicillin-streptomycin (60 U/mL). *cpnD* mutants were cultured in media containing blasticidin (30 µg/mL, InvivoGen, ant-bl-1). The *cpnD* mutant cell lines were generated via restriction enzyme-mediated integration (REMI) and are part of the genome-wide *Dictyostelium* insertion (GWDI) bank [33]. *cpnD* mutant cell lines consist of a 1,589 bp REMI insert containing a blasticidin-resistant gene (*bsr*) inserted into the first (291 bp) or second (459 bp) exon of the endogenous *cpnD* gene to create the *cpnD(i291)* and *cpnD(i459)* cells, respectively. The *cpnD* mutant cell lines were clonally isolated, and the location of the insertion was verified by PCR and sequencing. The cDNA for *cpnD* was obtained using RT-PCR and subcloned into the *SacI* site of the pTX-GFP plasmid [34,35]. The plasmid was electroporated into the AX4 parental cell strain. Cells were pulsed twice with a 5-second interval at 0.85 kV. Cells were placed on ice for 5 minutes and then plated in VL-6 media. Transformants were selected using G418 (geneticin) at 60 µg/mL after 24 hours.

### 2.2. Growth assay

AX4 and *cpnD* mutant cell lines were harvested from plates and placed in 125 mL flasks containing 12.5 mL of VL-6 media with 60 U/mL penicillin-streptomycin. Flasks containing a starting concentration of 5 x 10^4^ cells/mL were placed in a shaking incubator set to 125 RPM at 18 °C. Every 24 hours for 7 days, three samples per cell line were removed from each flask, and cell densities were estimated using a hemocytometer. The cell densities of the three samples from each flask were averaged. Three growth assay trials were performed, and the mean cell densities on each day were averaged. To determine significant differences between cell growth in AX4 and *cpnD* mutant cells, a repeated measures ANOVA with Tukey’s post hoc multiple comparisons test was performed.

### 2.3. Developmental assays

AX4 and *cpnD* mutant cell lines were harvested from plates, counted using a hemocytometer, and centrifuged at 437 x g for 5 minutes at 4 °C. Cell pellets were washed twice with ice-cold developmental buffer ((DB), 5 mM Na_2_HPO_4_, 5 mM KH_2_PO_4_, 1 mM CaCl_2_, 2 mM MgCl_2_, pH 6.5), and resuspended to obtain 5 x 10^7^ cells/mL. Cells (2.5 x 10^7^) were plated on black filters (catalog no. HABP04700; Millipore), placed on DB-soaked pads in dishes (catalog no. 09-753-53C; Fisher), and allowed to develop for 48 hours. Images were taken using a Nikon SMZ800N dissecting microscope every two hours for 28 hours, followed by a 48-hour image to observe mature fruiting bodies. Two developmental assays were performed, and representative images for each cell line were chosen. The area of the spore heads from each cell line (n > 700) across two trials was measured using ImageJ. Individual fruiting body measurements were graphed for each cell line after removing outliers. A one-way ANOVA with Tukey’s post hoc multiple comparisons test was performed to determine statistical significance between the AX4 and *cpnD* mutant cell lines.

For developmental assays on bacterial lawns, *E. coli B/R* was inoculated into 2 mL of VL-6 without penicillin-streptomycin and incubated at 220 RPM at 37 °C for 16 hours. AX4 and *cpnD* mutant cell lines were harvested from Petri dishes and counted using a hemocytometer and subjected to serial dilutions to obtain a final concentration of 500 cells/mL. *E. coli B/R* (500 µL) and *Dictyostelium* cells (100 µL) were spread on SM/5 agar plates (0.2% proteose peptone 2, 0.2% yeast extract, 0.2% glucose, 0.2% MgSO_4_×7 H_2_O, 0.19% KH_2_PO_4_, 1% K_2_HPO_4_, 1.5% agar, pH 6.4). Plates were incubated at 18 °C and examined for plaque formation. Plates were imaged with a Nikon SMZ800N dissecting microscope on days 3, 4, and 5.

### 2.4. Cell size measurements

AX4 and *cpnD* mutant cells were harvested from plates, counted using a hemocytometer, and centrifuged at 437 x g for 5 minutes at 4 °C. The cell pellet was resuspended in VL-6 media at 2 x 10^6^ cells/mL. On glass-bottom dishes, 200 µL of the cell suspension was plated and allowed to adhere for 20 minutes. Cells were then imaged using a TE2000-S Nikon Eclipse microscope with a 60X oil objective and differential interference contrast (DIC) microscopy. The area of individual cells from each cell type (n > 700) across two trials was measured using ImageJ, and individual cell measurements were graphed for each cell type after removing outliers. A one-way ANOVA with Tukey’s post hoc multiple comparisons test was performed to determine statistical significance between the AX4 and *cpnD* mutant cell lines.

### 2.5. Adhesion assay

AX4 and *cpnD* mutant cells were harvested from plates, counted using a hemocytometer, and centrifuged at 437 x g for 5 minutes at 4 °C. The cell pellets were resuspended in VL-6 media at 1 x 10^6^ cells/mL. On 60 mm Petri dishes, 2 mL of each cell type was allowed to adhere for 30 minutes. The cells that had not adhered to the plates were removed, and the media was replaced with fresh media. Images of the adhered cells were taken on a Nikon TE2000 microscope with a 20X phase-contrast objective at three marked spots on the Petri dishes. Cells on plates were rotated for 15 minutes at 50 RPM. Detached cells were removed from the plates, and new media was added. New images at each marked spot were captured. This procedure was repeated at 75 and 100 RPM. The number of cells in each image was counted using the ImageJ Cell Counter plugin, and counts were averaged. The percentage of detached cells after each rotation was calculated by the difference of the average number of cells remaining after rotation from the average number of cells present before rotation, divided by the number of cells present before rotation. Data from five trials were averaged and analyzed for significant differences between AX4 and *cpnD* mutant cells by a two-way ANOVA with uncorrected Fisher’s LSD post hoc test.

### 2.6. Ras activation assay

AX4 and *cpnD* mutant cells were harvested from plates, counted using a hemocytometer, and centrifuged at 437 x g for 5 minutes at 4 °C. The cell pellets were resuspended in VL-6 media at 4 x 10^6^ cells/mL. To assess Ras activity in the AX4 and *cpnD* mutant cells, we used an activated Ras pull-down assay kit (#BK008-S, Cytoskeleton Inc) following the manufacturer’s protocol, with the exception that 4 x 10^6^ cells/mL were used for the pull-down assays. Total Ras whole cell samples and activated Ras samples were analyzed by Western blot with an antibody to Ras (see section 2.7). Data from three and four trials were averaged for the activated Ras and total Ras, respectively. The data were analyzed for significant differences by a one-way ANOVA with Tukey’s post hoc multiple comparisons test.

### 2.7. Western blot

AX4 and *cpnD* mutant cells were harvested from plates, counted using a hemocytometer, and centrifuged at 437 x g for 5 minutes at 4 °C. The cell pellets were resuspended in sample buffer (0.2 M Tris-HCl, 0.4 M DTT, 277 mM SDS, 6 mM Bromophenol blue, 4.3 M Glycerol), and 2 x 10^6^ cells were loaded into the stain-free gels (BIO-RAD, 4568034) and run for 2 hours at 100 volts. After electrophoresis, the stain-free gels were imaged using a BIO-RAD ChemiDoc Touch Imaging System. The stain-free gels were then transferred to a polyvinylidene difluoride (PVDF) membrane (ThermoFisher Scientific, 88518) for 1 hour at 100 volts. The membranes were placed in Blotto (5% dried milk in 1% (v/v) Tween-20 in phosphate-buffered saline (PBS-T)) for 30 minutes, and then incubated with a-RegA (1:4000) polyclonal antibody (gift from Robert Kay; [36]), a-SibA (1:1000) polyclonal antibody (Geneva Antibody Facility) for 2 hours at room temperature, or a-Pan Ras (1:250) mouse monoclonal primary antibody for Ras pulldown samples (#BK008-S, Cytoskeleton) for 3 hours at room temperature. The membranes were washed three times for 5 minutes with PBS-T. After washing, the membranes were incubated with an a-rabbit HRP-conjugated secondary antibody (1:15,000) or an a-mouse HRP-conjugated secondary antibody (1:15,000) for 1 hour at room temperature in Blotto. The membranes were washed three times for 10 minutes with PBS-T. Membranes were imaged on the BIO-RAD ChemiDoc Touch Imaging system using Femtoglow HRP substrate (Michigan Diagnostics, FWPS02).

### 2.8. Western blot densitometry

For densitometry analysis of SibA, band densities were measured using Image Lab v6.1 from BIO-RAD. Each band density of the Western blot image was measured and then normalized to the density of all proteins in the corresponding lane from the stain-free gel image. Normalized Western blot band densities were further normalized to the average adjusted total band volume intensities across all lanes. Western blot and gel images from six trials were analyzed, and normalized band densities were averaged and analyzed for significant differences using a one-way ANOVA with Tukey’s post hoc multiple comparisons test.

For densitometry analysis of total Ras, each band density of the Western blot image was measured and then normalized to the density of all proteins in the corresponding lane from the stain-free gel image. Normalized Western blot band densities were further normalized to the average adjusted total band volume intensities across all lanes. Western blot and gel images from four trials were analyzed, and normalized band densities were averaged and analyzed for significant differences using a one-way ANOVA with Tukey’s post hoc multiple comparisons test.

For densitometry analysis of active Ras, band densities of active Ras and total Ras were measured using Image Lab v6.1 from BIO-RAD. The adjusted total band volume densities of the active Ras bands were normalized to the average adjusted total band volume intensities across all active Ras lanes, and the same was done for total Ras as described above. Normalized active Ras band densities were further normalized to the normalized adjusted total band volume intensities of their respective total Ras samples. Western blot images from three trials were analyzed, and normalized band densities were averaged and analyzed for significant differences using a one-way ANOVA with Tukey’s post hoc multiple comparisons test.

### 2.9. Contractile vacuole assay

AX4 and *cpnD* mutant cells in VL-6 media (2 x 10^6^ cells/mL) were plated on 35 mm glass-bottom dishes and allowed to adhere for 20 minutes. The media was removed and replaced with water or water with 25 µM LY294002 (Cell Signaling Technologies, #9901S). After a two-hour incubation period, the cells were imaged using a TE2000-S Nikon Eclipse microscope with a 60X oil objective and differential interference contrast (DIC) microscopy. The area of each contractile vacuole (n > 209 CVs/cell type) was measured using ImageJ, outliers were removed, and a one-way ANOVA with Šídák’s post hoc multiple comparisons test was performed to analyze significant differences between AX4 and each of the *cpnD* mutant cells, as well as differences between cell types and LY294002 treatment.

### 2.10. GFP-tagged CpnD assays

Parental cells containing the GFP-CpnD expression plasmid were harvested from plates, counted using a hemocytometer, and centrifuged at 437 x g for 5 minutes. The cells were resuspended in DB at 5 x 10^6^ cells/mL. In 35 mm glass-bottom dishes, 2 mL of cell suspension was plated and allowed to starve for 8 hours. After 8 hours, the DB was removed, and 200 µL of fresh DB with 2 mM caffeine was added to the central well of the glass-bottom dish and allowed to incubate for 30 minutes. After incubation, 100 µL of the cell suspension was removed from the plates before confocal time-lapse imaging began (images taken every 3 seconds). After 10 seconds of imaging, 100 µL of 10 µM cAMP in DB was added. Images were obtained using a Nikon A1R confocal microscope with a 60x oil objective. Images were processed in Adobe Photoshop, and levels of brightness and contrast were adjusted. The membrane on and off times were estimated from 7 cells across 3 trials, and average times were calculated.

## 3. Results

### 3.1 cpnD mutants have increased proliferation in both axenic and bacterial cultures

To study the function of CpnD in *Dictyostelium*, we obtained two *cpnD* mutant cell lines and the parental (AX4) strain from the *Dictyostelium* Stock Center [32]. The *cpnD* mutants were created as part of a REMI-seq project to generate a genome-wide mutant resource for *Dictyostelium* [33]. The *cpnD* mutants contain a 1,589 bp insert in the first (*cpnD(i291)*) or second (*cpnD(i459)*) exon of the endogenous *cpnD* gene, respectively (Figure 1A). We isolated clonal populations of each cell strain and confirmed the presence and location of the REMI insertions in each mutant via PCR (Supplemental Figure 1) and Sanger sequencing.

**Figure 1.**
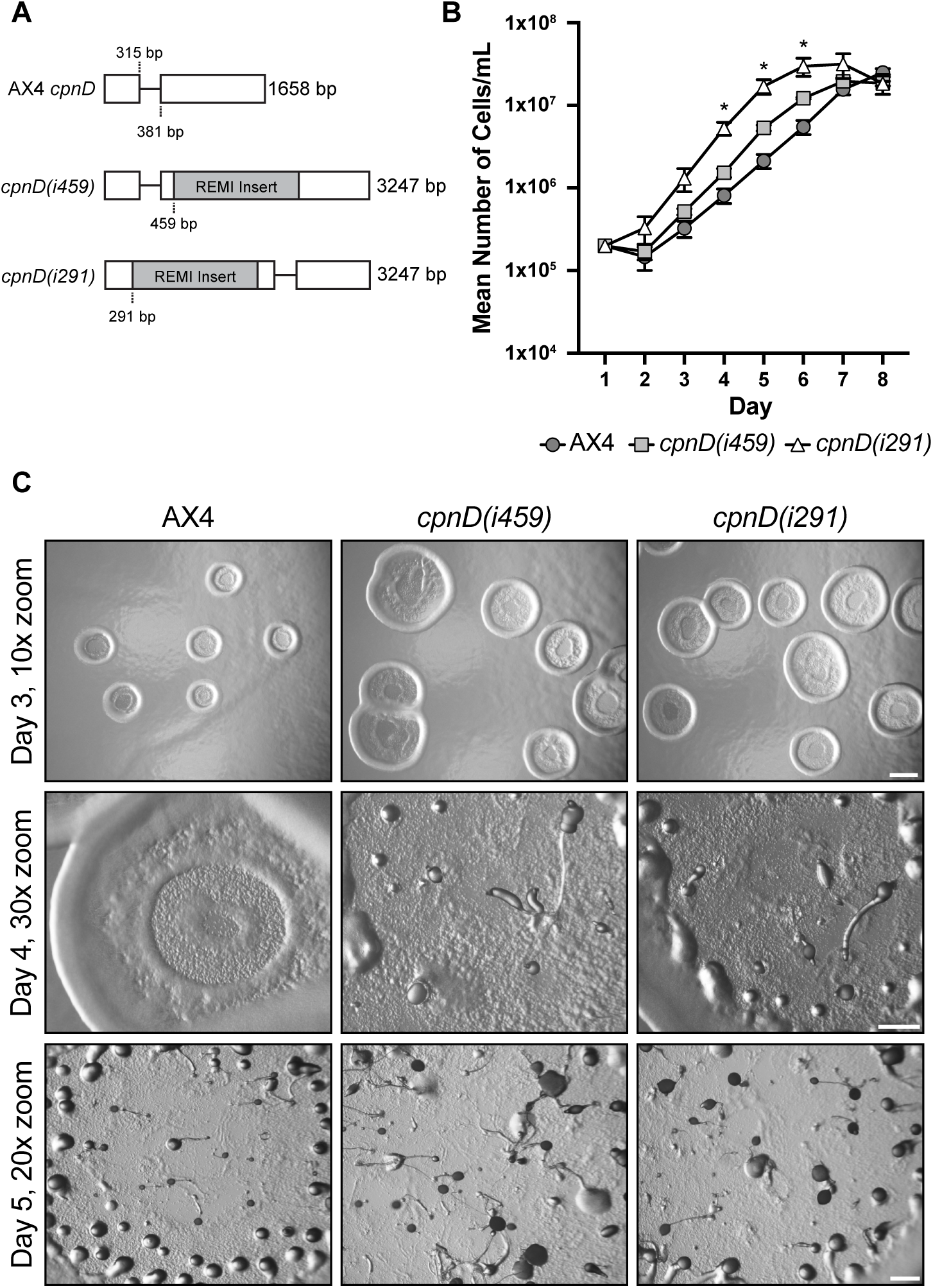
*cpnD* mutants have increased proliferation in both axenic and bacterial cultures. (**A**) Schematic showing the location of the 1,589 bp REMI insert in the first (*cpnD(i291))* or second (*cpnDi(459))* exon of the endogenous AX4 *cpnD* gene in the two *cpnD* mutants. (**B**) AX4 cells and both *cpnD* mutants were grown in a shaking suspension starting at a cellular density of 5 x 10^4^ cells/mL. Cell densities were counted and averaged once a day for 7 days using a hemocytometer. Mean cell densities were calculated from three trials, and a repeated measures ANOVA with Tukey’s post hoc multiple comparisons test was performed to analyze significant differences between cell lines. * indicates a significant difference between the parental AX4 cell line and *cpnD(i291)*, *p* < 0.05. Error bars = standard error. (**C**) AX4 and *cpnD* mutant cells were plated with *E. coli B/R* on SM/5 agar. Plaques were imaged after 3, 4, and 5 days after plating, and representative images are shown. Day 3 scale bar = 1000 µm. Day 4 and 5 scale bars = 500 µm.

To determine if the mutants exhibited normal proliferation, we performed growth assays with both the parental and *cpnD* mutant cell lines. Cells were grown in a shaking suspension, and cell samples were collected and counted using a hemocytometer each day for 7 days. Both *cpnD(i291)* and *cpnD(i459)* cells appeared to proliferate faster than the parental cell line, but only the *cpnD(i291)* cells were significantly different from the parental AX4 cells (Figure 1B). We also examined cell proliferation on bacterial lawns. AX4 cells and *cpnD* mutants were plated on agar with *E. coli B/R*. As *Dictyostelium* feeds on the *E. coli B/R*, clear plaques arise within the bacterial lawn. Images were taken of the plaques on days 3, 4, and 5 after plating. The plaques created by the *cpnD* mutants appeared on day 2, while AX4 plaques appeared on day 3. *cpnD* mutant plaques were also noticeably larger than the parental AX4 plaque on day 3 (Figure 1C). As cells consume the bacteria at the edge of the plaque, cells within the plaque starve and go through development to form fruiting bodies. On day 4, both *cpnD* mutants had begun development with the formation of mounds and slugs, while AX4 cells had not yet formed multicellular structures. On day 5, AX4 cells had formed mounds and a few fruiting bodies, while the *cpnD* mutants had formed mostly fruiting bodies (Figure 1C). These data indicate that *cpnD* mutants proliferate faster in both axenic and bacterial cultures.

### 3.2. cpnD mutants exhibit precocious development and have larger fruiting bodies than parental cells

We next investigated whether *cpnD* mutants exhibited normal developmental morphology and timing by monitoring the synchronized development of both the *cpnD* mutants and parental AX4 cells. Parental and *cpnD* mutants in starvation buffer were plated on nitrocellulose filters, and images were captured throughout a 48-hour timeframe. Representative images of all three cell lines are shown at 4 different timepoints in Figure 2A. At 6 hr, none of the three cell lines displayed multicellular structures. At 18 hr, AX4 cells had formed small mounds. In contrast, *cpnD(i459)* cells had progressed beyond the mound stage and formed large slugs, while the *cpnD(i291)* cells were at an intermediate stage of development, forming large mounds with some beginning to tip and transition into slugs. These observations indicate that both *cpnD* mutants develop more rapidly than AX4 cells, with *cpnD(i459)* cells developing faster than *cpnD(i291)* cells. Additionally, both *cpnD* mutants appeared to form larger mounds, slugs, and fruiting bodies compared to AX4 cells (Figure 2A). To quantify these differences, we measured the area of fruiting body spore heads (n > 700 across two trials) using ImageJ. Both *cpnD* mutants produced significantly larger spore heads than AX4, with *cpnD(i459)* forming significantly larger spore heads than *cpnD(i291)* (Figure 2B).

**Figure 2.**
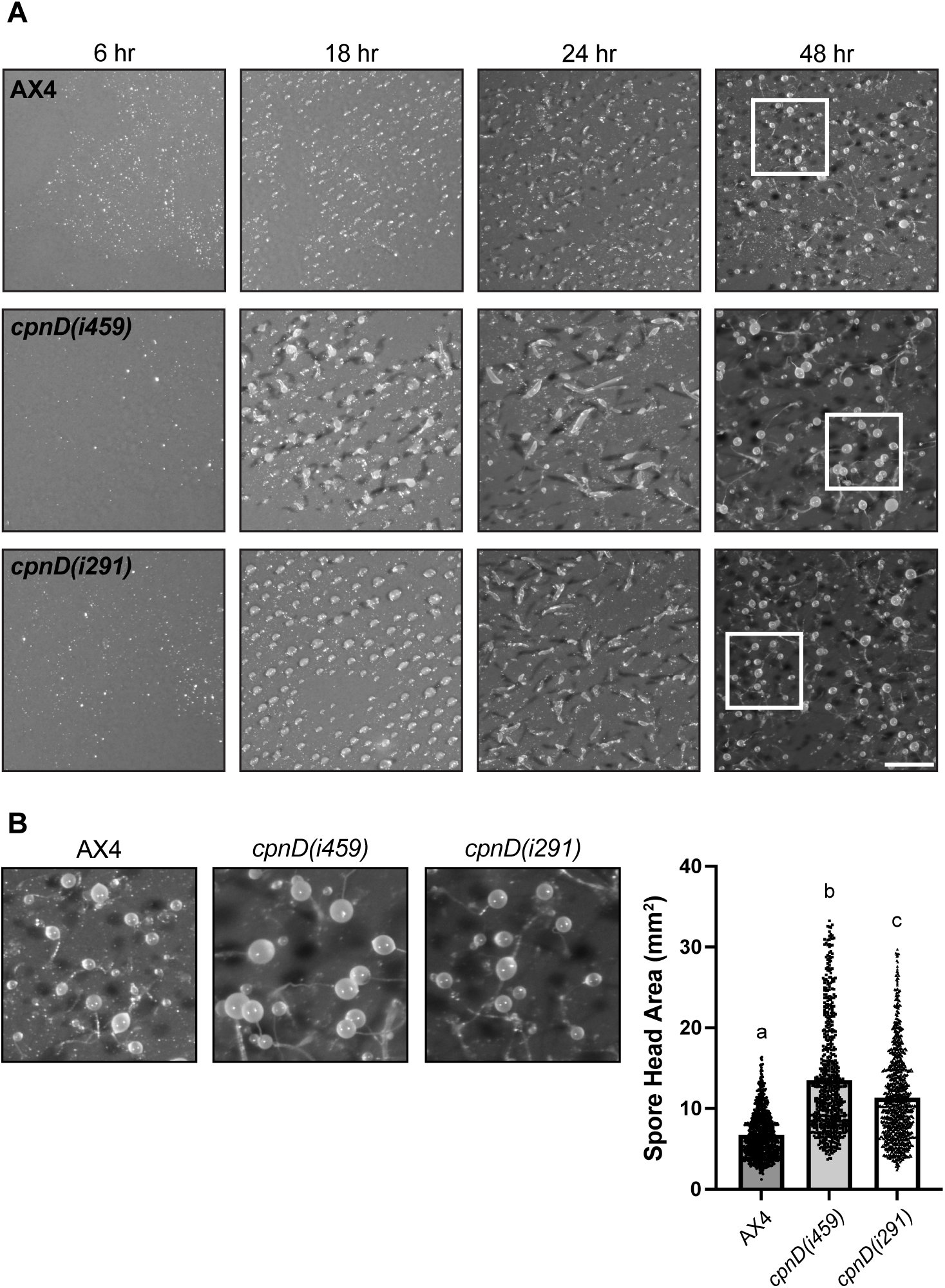
*cpnD* mutants exhibit precocious development and have significantly larger fruiting bodies. (**A**) AX4 and *cpnD* mutant cells were resuspended in starvation buffer and plated on nitrocellulose filters to synchronize developmental timing. Representative images across two trials are shown throughout a 48-hour imaging window. At 6 hr, none of the cell lines exhibited multicellular characteristics. At 18 hr, AX4 cells formed small mounds, while *cpnD(i459)* mutants formed large slugs and *cpnD(i291)* formed large mounds with some slugs. At 24 hr, AX4 cells formed slugs, while both *cpnD* mutants formed larger slugs and showed occurrences of fruiting body formation. At 48 hr, all three cell lines reached their terminal developmental timepoint, exhibiting fruiting bodies of various sizes. Scale bar = 1 mm. (**B**) Insets from white boxes shown in Figure 2A of AX4 and *cpnD* mutant cells at 48 hr developmental time point. The bar graph represents the area of fruiting body spore heads (n > 700 across two trials) measured using ImageJ. Data were analyzed for significant differences using a one-way ANOVA with Tukey’s post hoc multiple comparisons after removing outliers. Different letters above the bars indicate a significant difference, *p* < 0.0001. Error bars = standard error.

### 3.3. cpnD mutant cells display a flattened morphology and are significantly larger than the parental cell line

Upon morphological inspection of the *cpnD* mutant cells, we noticed that some of the mutant cells appeared flatter and “pancake”-shaped compared to the parental AX4 cell line. To quantify these observations, we imaged cells using differential interference contrast (DIC) microscopy and measured the cell area. On average, both the *cpnD* mutants had a significantly larger mean cell area than the parental cell line. Interestingly, *cpnD(i459)* cells were the largest cell type and significantly differed in size from the *cpnD(i291)* cells (Figure 3A, 3B).

**Figure 3.**
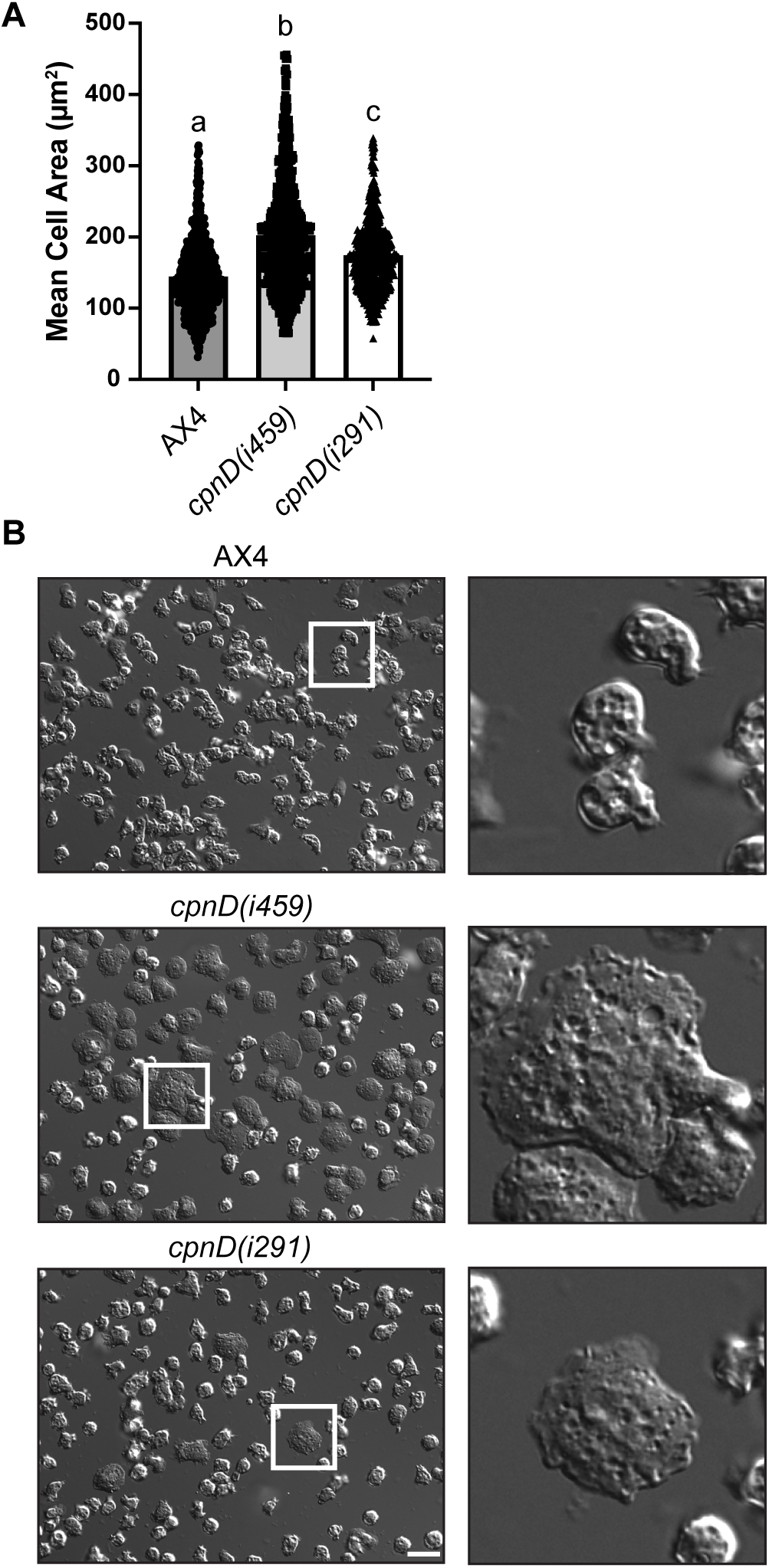
*cpnD* mutants have a larger cell area than parental cells. (**A**) The mean cell area of AX4 and *cpnD* mutants (n > 700 cells across two trials) was measured using ImageJ, and the bar graph represents individual cell measurements. Data were analyzed for significant differences using a one-way ANOVA with Tukey’s post hoc multiple comparisons after removing outliers. Different letters above the bars indicate a significant difference, *p* < 0.0001. Error bars = standard error. (**B**) AX4 and *cpnD* mutants were plated on glass-bottom dishes and imaged using DIC microscopy. Representative images are shown with insets from white boxes. Scale bar = 25 µm.

### 3.4. cpnD mutants have decreased cell-substrate adhesion

The flatter, pancake-like morphology of the *cpnD* mutant cells suggested that *cpnD* mutants may have altered substratum adhesion; increased and decreased adhesion have both been shown to be associated with a flattened cellular morphology [37,38]. To test this, parental and *cpnD* mutants were allowed to adhere to Petri dishes, then rotated at 3 speeds (50, 75, and 100 RPM). After each rotation, the media was replaced to remove detached cells. Three marked locations on the dish were imaged using phase contrast microscopy to assess the number of cells that remained adhered to the dish. We found that both *cpnD* mutants showed significantly reduced adhesion to the Petri dish at 50 and 75 RPM compared to the parental cell line (Figure 4A). There were no significant differences in cell-substrate adhesion between the parental and *cpnD* mutant cell lines at 100 RPM, suggesting that the forces inflicted on the cells at high RPM are greater than the adhesion strength of *Dictyostelium*.

**Figure 4.**
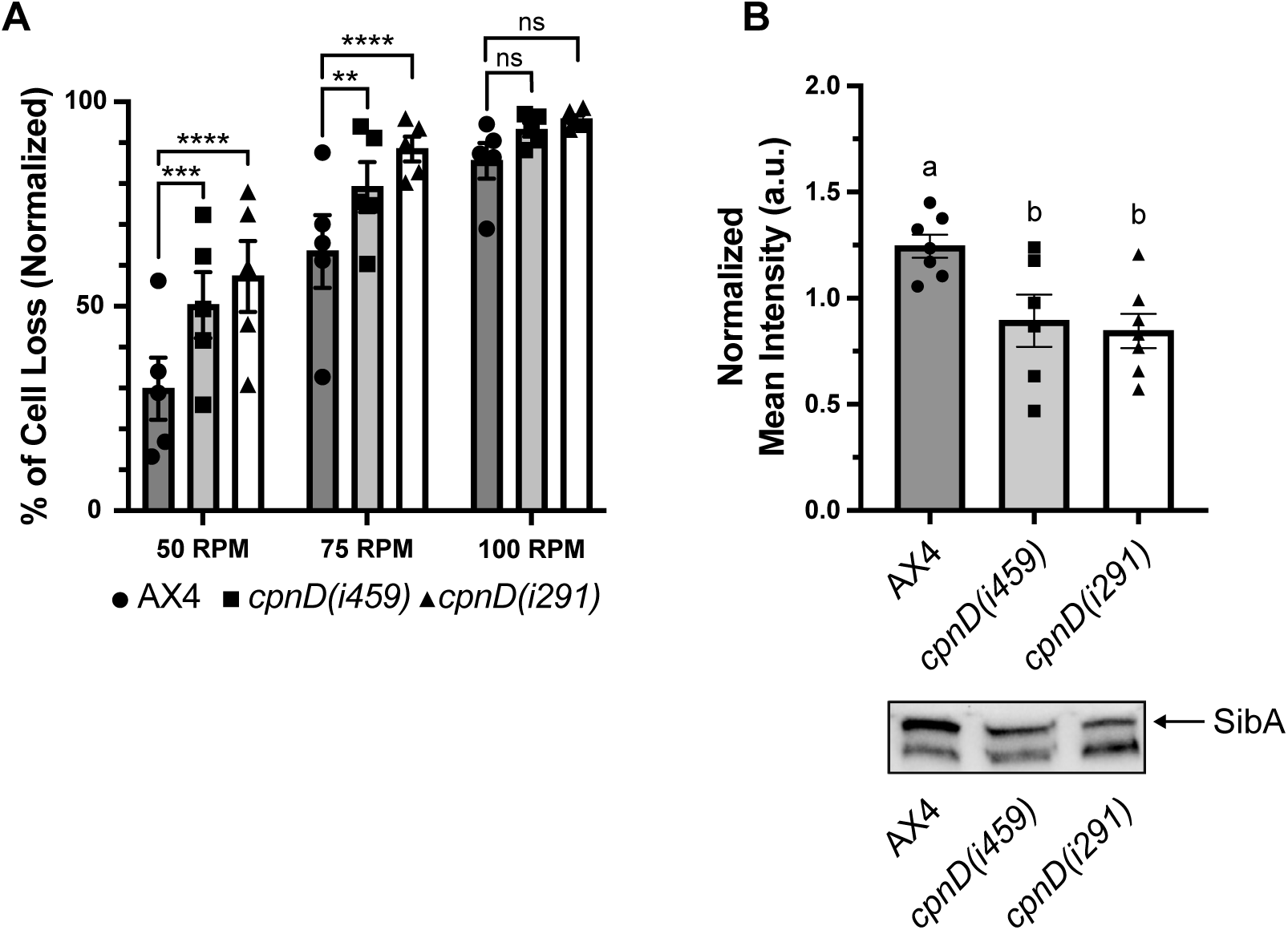
*cpnD* mutants have reduced cell-substrate adhesion and SibA expression. (**A**) AX4 and *cpnD* mutant cell lines were plated on 60 mm Petri dishes. Cells were imaged with phase-contrast microscopy at three marked spots before and after rotation at 50, 75, and 100 RPM. The number of cells in each image was counted and averaged. The percentage of detached cells after rotation was calculated by subtracting the average number of cells remaining after each rotation from the average number of cells present before rotation, divided by the number of cells present before rotation. Data from five trials were averaged and analyzed for significant differences between the AX4 and *cpnD* mutant cell lines using a repeated measures two-way ANOVA with uncorrected Fisher’s LSD post hoc test. ** indicates *p* < 0.01, *** indicates *p* < 0.001, **** indicates *p* < 0.0001, ns indicates no significant difference. Error bars = standard error. (**B**) Densitometry analysis of the SibA band on Western blots. The mean intensity of the SibA Western blot band from AX4 and *cpnD* mutants was normalized to their respective lanes on stain-free gel images. The bar graph represents the average normalized band densities from 6 trials, and these data were analyzed for significant differences using a one-way ANOVA with Tukey’s post hoc multiple comparisons test. Different letters indicate a significant difference (*p* < 0.05) while the same letters indicate no significant difference. A representative Western blot analysis of SibA expression in AX4 and *cpnD* mutant cells is included below the bar graph. Error bars = standard error.

SibA is a transmembrane protein essential for cell-substrate adhesion in *Dictyostelium* and has been shown to affect cellular adhesion based on its expression [39]. We have previously shown that *cpnC-* cells have significantly decreased cell-substrate adhesion and reduced SibA expression [31]. To determine if the decreased cell-substrate adhesion in *cpnD* mutants may be due to changes in SibA expression, whole-cell samples of parental and *cpnD* mutant cells were analyzed by Western blot to determine levels of SibA expression. SibA expression was reduced in both *cpnD* mutants compared to parental cells (Figure 4B). Because the a-SibA antibody recognizes an additional protein, we have previously performed Western blotting with *sibA* null cells to verify that the higher molecular weight band is SibA [31].

### 3.5. cpnD mutant cells have increased levels of activated Ras

The flatter, pancake-like cellular morphology has been observed in cells that have increased activation of RasC [21], possibly due to the activation of the actin cytoskeleton, causing cells to create protrusions and spread out [40]. Additionally, research has shown that global activation of Ras by recruitment of GefA to the cellular membrane promotes cell spreading [41]. Because our *cpnD* mutants were significantly larger and flatter than the parental cell line, we wanted to test whether the *cpnD* mutants had increased activated Ras. To test this, we performed an activated Ras pulldown assay. Normally, vegetative cells that have not been starved and have not been stimulated with cAMP show very little Ras activation [14]. However, our vegetative *cpnD* mutants did have significantly higher levels of active Ras compared to the parental strain, while the levels of total Ras between all cell lines were not significantly different (Figure 5).

**Figure 5.**
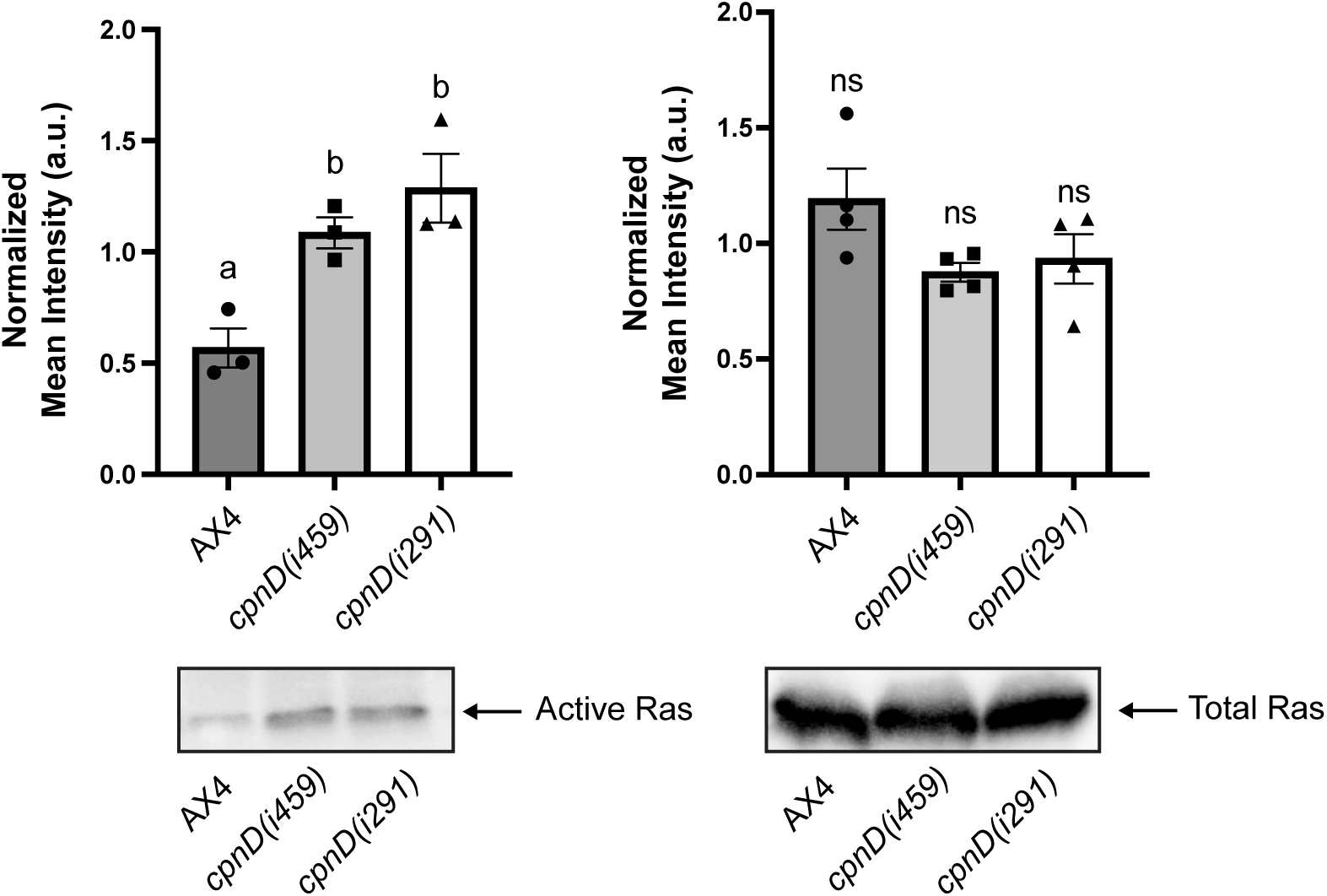
*cpnD* mutants have increased levels of activated Ras. Densitometry analysis of the activated and total Ras bands on Western blots. The mean intensity of total Ras alone was measured and normalized to the density of all proteins in the corresponding lane from the stain-free gel image. Normalized Western blot band densities were further normalized to the average adjusted total band volume intensities across all lanes. Data were analyzed across four trials, and significant differences were determined using a one-way ANOVA with Tukey’s post hoc multiple comparisons test. ns indicates no significant difference. The adjusted total band volume densities of the activated Ras Western blot bands from AX4 and *cpnD* mutants were normalized to the average adjusted total band volume intensities across all active Ras lanes. Normalized active Ras band densities were further normalized to the normalized adjusted total band volume intensities of their respective total Ras samples. Data were analyzed across three trials, and significant differences were determined using a one-way ANOVA with Tukey’s post hoc multiple comparisons test. Different letters above the bars indicate significant differences (*p* < 0.05). Representative Western blot analyses of active Ras and total Ras are shown below their respective bar graphs. Error bars = standard error.

### 3.6. cpnD mutants have a reduced CV area

Contractile vacuoles are intracellular organelles involved in the regulation of water and various ions in numerous cells, including *Dictyostelium* [42]. We have previously shown that *cpnA-* cells have larger and more persistent CVs [26,28]. Because of the role CpnA plays in CV function, we wanted to determine if *cpnD* mutants exhibited similar CV defects. To test this, we incubated parental and *cpnD* mutant cells on glass-bottom dishes in water for 2 hours before performing DIC microscopy and measuring CV area (n > 1,450 CVs per cell line across three trials). We found that *cpnD* mutants exhibited significantly smaller CVs compared to the parental cell line (Figure 6A,B). Interestingly, the *cpnD(i459)* mutants had the smallest average CV area, while the average CV area of *cpnD(i291)* cells was significantly larger than *cpnD(i459)* mutants, but still significantly smaller than the parental cells.

**Figure 6.**
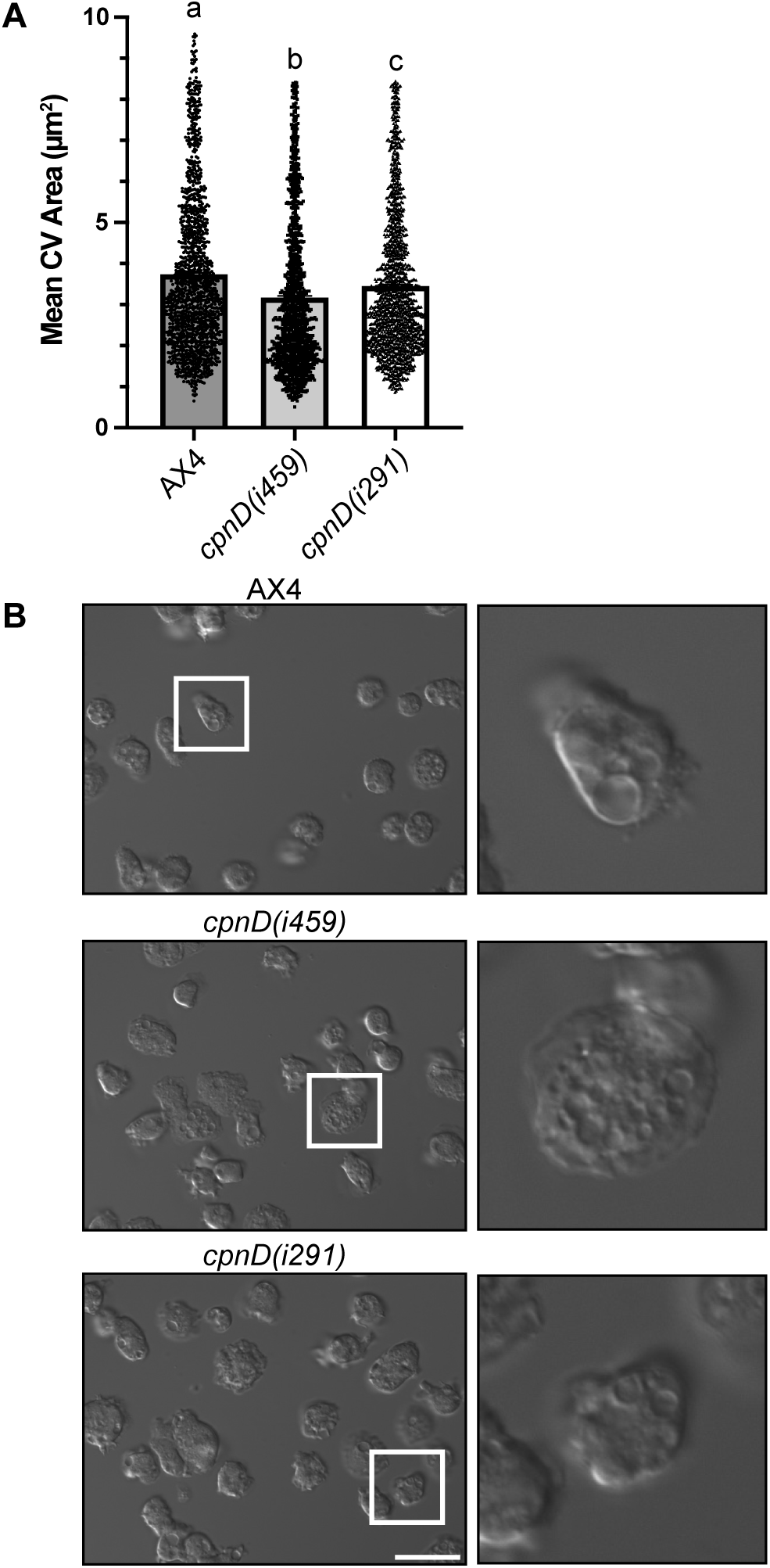
*cpnD* mutant cells have a reduced CV area. AX4 and *cpnD* mutants were plated on glass-bottom dishes and incubated in water for two hours and then imaged using DIC microscopy. (**A**) The mean CV area of AX4 and *cpnD* mutants (n > 1450 cells across four trials) was measured using ImageJ, and the bar graph represents individual cell measurements. Data were analyzed for significant differences using a one-way ANOVA with Tukey’s post hoc multiple comparisons after removing outliers. Different letters above the bars indicate a significant difference, *p* < 0.0001. Error bars = standard error. (**B**). Representative images are shown with insets from white boxes. Scale bar = 25 µm.

### 3.7. PI3K inhibition suppresses the large cell size and small CV defects observed in cpnD mutants

To determine if the increased Ras activation observed in *cpnD* mutants is responsible for their flatter morphology, we used the PI3K inhibitor LY294002 to inhibit PI3K, which is activated by Ras. We performed DIC microscopy to image cells in the absence or presence of LY294002 treatment (n > 180 cells per cell line across two trials) and measured the mean cell area. As we previously observed (Figure 3), the mean cell area of the *cpnD* mutants was significantly larger than that of the parental cell line. However, upon LY294002 treatment, the mean cell area of both *cpnD* mutant cell lines was significantly reduced, so much so that the *cpnD(i291)* cells were no longer significantly larger than the parental cell line (Figure 7A).

**Figure 7.**
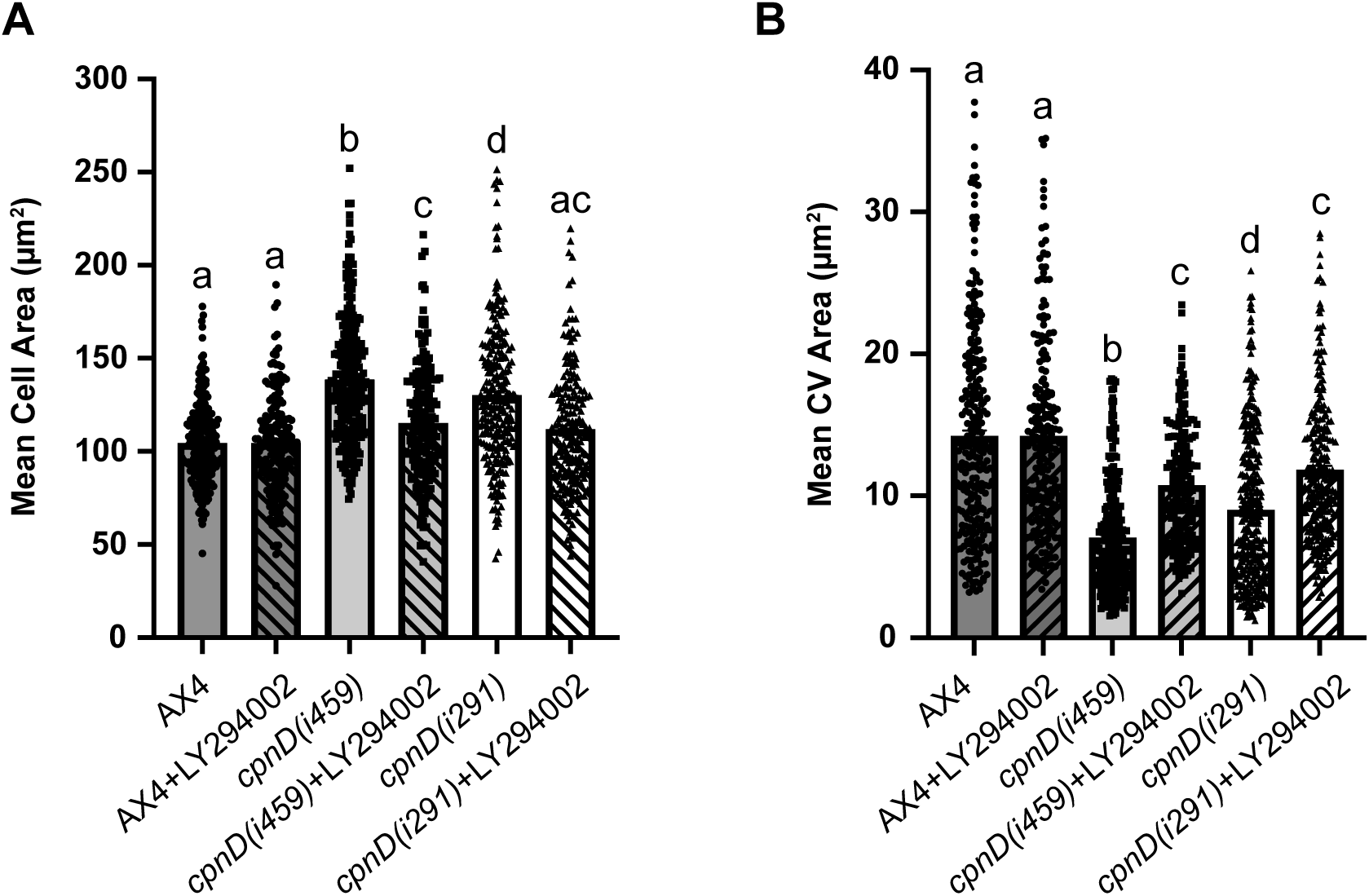
*PI3K inhibition suppresses both cell and CV size defects.* (**A**) AX4 and *cpnD* mutant cells were plated on glass-bottom dishes in the absence or presence of 25 µM PI3K inhibitor (LY294002) and imaged with DIC microscopy. The mean cell area of AX4 and *cpnD* mutants (n > 180 cells across two trials) was measured using ImageJ, and the bar graph represents individual cell measurements. Data were analyzed for significant differences using a one-way ANOVA with Tukey’s post hoc multiple comparisons after removing outliers. Different letters above the bars indicate a significant difference, *p* < 0.0001. Error bars = standard error. (**B**) AX4 and *cpnD* mutant cells were plated on glass-bottom dishes and incubated in water in the absence or presence of 25 µM PI3K inhibitor. The mean area of CVs (n > 209 CVs per cell line) was measured using ImageJ. After removing outliers, the data across two trials were analyzed for significant differences using a one-way ANOVA with Tukey’s post hoc multiple comparisons test. Different letters above the bars indicate a significant difference, *p* < 0.0001. Error bars = standard error.

To determine whether the small CV phenotype observed in *cpnD* mutants was associated with Ras/PI3K overactivation, we searched for similar phenotypes reported in the literature. One study identified *sodC*, which encodes a superoxide dismutase, in a screen for mutants with constitutively active PI3K [43]. Further studies showed that *sodC*- cells exhibited smaller than normal CVs that could be rescued by the PI3K inhibitor, LY294002 [44]. To determine whether LY294002 could suppress the small CV defect observed in the *cpnD* mutants, we performed CV assays with LY294002-treated and untreated parental AX4 and *cpnD* mutant cell lines. Cells were allowed to adhere to glass-bottom dishes and then incubated in water with or without LY294002 before performing DIC microscopy to measure the individual area of CVs (n > 200 CVs per cell line across two trials). As we previously observed, the mean CV area of the *cpnD* mutants was smaller compared to AX4 (Figure 7B). However, when we treated the cells with LY294002, the mean CV area of the *cpnD* mutant cell lines significantly increased, while the CV area of the parental AX4 strain did not change (Figure 7B). These experiments indicate that the flat morphology and small CV defects observed in *cpnD* mutants are due to increased activation of the Ras/PI3K signaling pathway.

### 3.8. GFP-tagged CpnD translocates from the cytosol to the plasma membrane in response to cAMP stimulation

Ras proteins are uniformly distributed along the plasma membrane of *Dictyostelium* cells, and it has been shown that cAMP stimulation results in the activation of Ras (Ras-GTP) from its inactive state (Ras-GDP) [45,46]. Because Ras and subsequently PI3K are activated at the plasma membrane in response to cAMP stimulation, Ras and PIP_3_ localize at the leading edge of chemotaxing cells [13,15,21,22], we hypothesize that if CpnD is involved in this pathway, CpnD will translocate to the plasma membrane in response to cAMP stimulation. To test this hypothesis, we subcloned the cDNA for *cpnD* into the pTX-GFP vector at the *SacI* site for the overexpression of CpnD tagged with GFP at the N-terminus (GFP-CpnD) in the parental cell line. Cells were starved in DB for 8 hours to induce aggregation and then incubated with caffeine for 30 minutes to suppress the release of endogenous cAMP. Time-lapse confocal microscopy was used to image cells before and after cAMP stimulation. In some cells, we captured GFP-tagged CpnD translocating from the cytosol to the plasma membrane and then returning to the cytosol (Figure 8, Supplemental Video 1). The average time of translocation after cAMP stimulation was 24 ± 4 seconds, while the average time on the membrane was 18 ± 4 seconds (n = 7 cells, across 3 trials). These experiments indicate that CpnD translocates to the plasma membrane in response to cAMP and could potentially play a role in the regulation of Ras at the plasma membrane.

**Figure 8.**
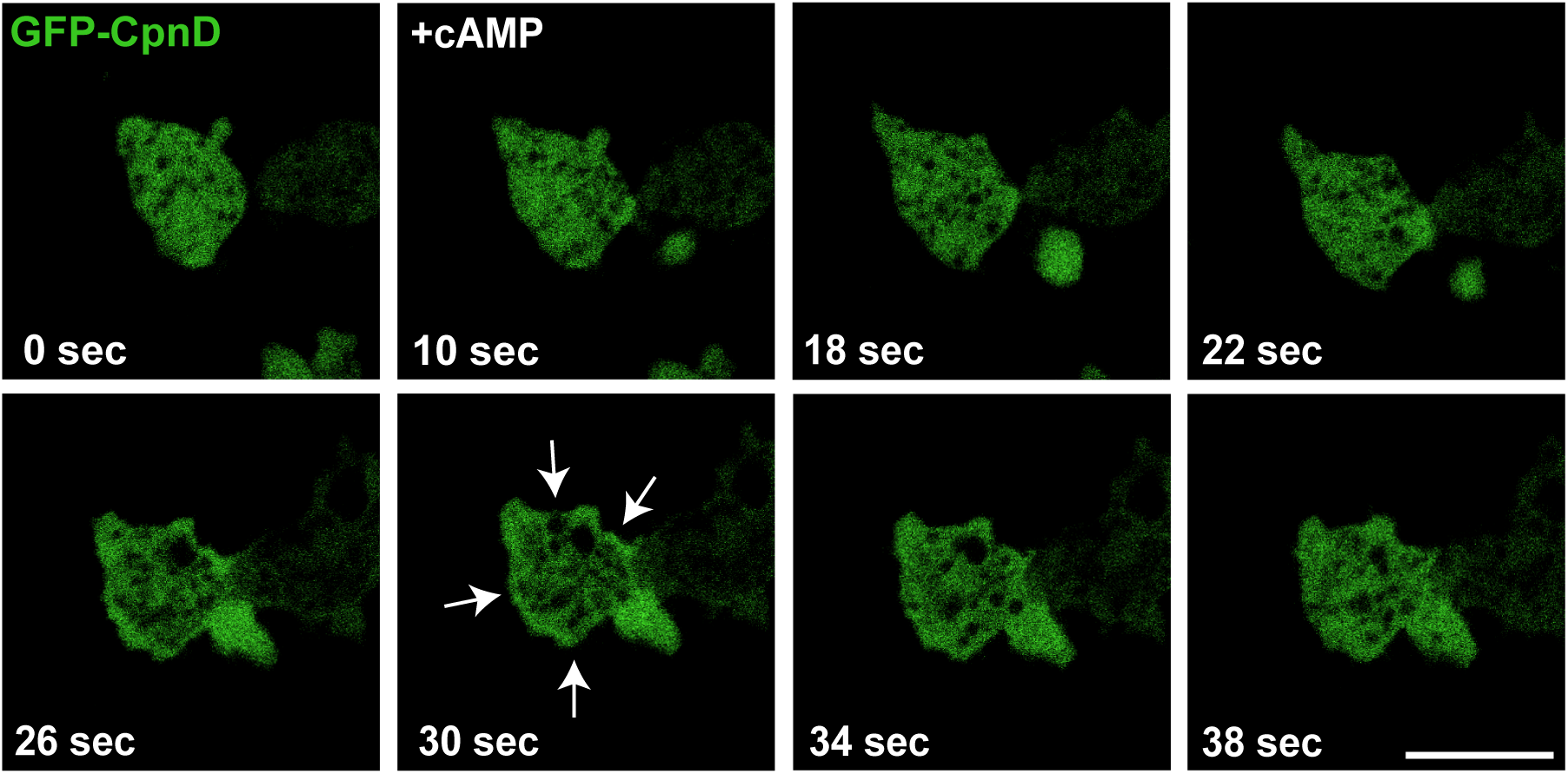
GFP-tagged CpnD translocates to the plasma membrane in response to cAMP stimulation. Parental cells expressing GFP-CpnD were resuspended in DB and plated on a 35-mm glass-bottom dish before an 8-hour starvation period. After 8 hours, the DB was removed from the dish, and the cells were incubated in 2 mM caffeine for 30 minutes. Following caffeine treatment, time-lapse confocal microscopy images were obtained every 3 seconds, and 5 µM cAMP was added after 10 seconds of imaging (+cAMP in figure). Images were obtained using a Nikon A1R confocal microscope with a 60x oil immersion objective. Representative images are shown. Arrows indicate GFP-CpnD at the cell membrane. Scale bar = 10 µm.

## 4. Discussion

To study the function of CpnD, we obtained two REMI mutants from the *Dictyostelium* Stock Center that differed in the location of the insert within the *cpnD* gene [33]. The *cpnD(i291)* cell line contains the REMI insert within the first exon of the endogenous *cpnD* gene, while the *cpnD(i459)* cell line contains the REMI insert in the second exon. We initially hypothesized that the phenotypic defects of the *cpnD(i459)* mutant with the REMI insert after the first C2 domain might be less severe because the allele may encode a partially functional protein. However, we observed the opposite; the phenotypic defects seen in the *cpnD(i459)* mutant were more severe than those of the *cpnD(i291)* mutant. One possible explanation is that the *cpnD(i459)* mutant is making a truncated CpnD protein containing the C2 domain that exerts a dominant-negative effect. The truncated protein may retain the ability to bind some target proteins but lack normal function, thereby blocking other proteins from accessing those same targets.

The Ras activation assays we performed did not allow us to identify which of the different Ras isoforms were overactivated in the *cpnD* mutants. However, both RasC and RasG have been shown to be involved in the activation of PI3 kinases that control early developmental cAMP signaling and various actin-mediated functions like chemotaxis, macropinocytosis, and phagocytosis [15,47,48]. We hypothesize that all the phenotypes observed in the *cpnD* mutants can be attributed to the overactivation of Ras. Ras activity is regulated by RasGEFs and RasGAPs [49], and the *Dictyostelium* genome has at least 25 putative RasGEFs [50] and 14 putative RasGAPs [41] and previous studies with mutations that cause the overactivation of Ras have resulted in similar phenotypes. For example, expression of constitutively active RasC in *pten*- cells leads to cell spreading and a flattened morphology [21]. Likewise, recruitment of RasGEF to the plasma membrane also causes cell spreading [41].

Loss of the RasGAP NF1 allows for axenic growth by increasing macropinocytosis and results in increased phagocytosis [51]. Similarly, loss of the RasGAP C2GAP2 results in increased macropinocytosis and phagocytosis, which leads to increased growth rate [52]. Although we did not directly assess macropinocytosis and phagocytosis in the *cpnD* mutants, the increased growth in both axenic culture and increased plaque size on bacterial lawns could be attributed to increased macropinocytosis and phagocytosis, respectively. Ras is also activated in response to increased superoxide species, and *Dictyostelium* lacking superoxide dismutase (*sodC-*) have increased activated RasG [43,53]. *sodC-* cells also have significantly smaller contractile vacuoles (CVs), a phenotype that could be suppressed upon inhibition of PI3K with LY294002 [44,53]. The smaller CV phenotype observed in the *cpnD* mutants could also be suppressed with LY294002 treatment, indicating that increased Ras activation in these mutants also causes the small CV phenotype. *cpnD* mutants also exhibited precocious development and reduced cell-substratum adhesion, as well as decreased SibA expression. These phenotypes are also likely due to the higher levels of activated Ras in the *cpnD* mutants. Ras is activated in response to cAMP signaling, which regulates early development and changes in gene expression during development. *sibA* is a gene that is downregulated during early *Dictyostelium* development [31]; therefore, we hypothesize that the increased Ras activation in *cpnD* mutants results in precocious development, which is dependent on early developmental gene expression changes, like reduced *sibA* expression.

The higher levels of active Ras suggest that CpnD acts as a negative regulator of Ras. CpnD could act directly on Ras and function like a RasGAP, or CpnD could indirectly regulate Ras, perhaps through the regulation of a Ras-GAP or Ras-GEF. If CpnD acts upstream of Ras in response to cAMP stimulation, then we would expect CpnD to be located at the plasma membrane or translocate to the plasma membrane in response to cAMP. We previously showed that the other 5 copines in *Dictyostelium* are cytosolic proteins that bind to specific acidic phospholipids in a calcium-dependent manner and that all five copines transiently translocate to the plasma membrane in response to cAMP stimulation. Each of the five copine proteins translocated from the cytosol to the plasma membrane with different timing and magnitude [35]. Here, we show that CpnD also translocates from the cytosol to the plasma membrane in response to cAMP stimulation. The timing is most similar to GFP-tagged CpnC data [35], with the GFP-tagged CpnD showing an average “ON” time from cAMP stimulation of 24 ± 4 seconds, and lasting around 18 ± 4 seconds total.

Previous work has demonstrated that Ras activation has three distinct phases, with maximal translocation from the cytoplasm to the plasma membrane occurring 6 seconds after cAMP stimulation, followed by symmetry breaking, and then confinement of Ras activity to the leading edge of the cell 20-30 seconds later [13]. Because our data suggest that GFP-CpnD is transiently recruited to the membrane after the initial activation of Ras, we suspect CpnD may function during the symmetry breaking or confinement stages of Ras activation, possibly to facilitate negative feedback mechanisms that restrict Ras signaling during cellular polarization. CpnD may be acting similarly to the RasGAP C2GAP1, which also has two C2 domains. C2GAP1 binds to both Ras and activated Gα2 and is important in the regulation of adaptation to cAMP signals [54,55].

This study is the first to describe copine proteins as having a regulatory function in Ras activation and its subsequent downstream signaling effects. In genetic cancer screenings, it has been shown that numerous copine genes are dysregulated [3]. Additionally, studies have shown that constitutively active Ras leads to downstream signaling pathway dysregulation, and mutations that cause constitutively active Ras are highly prevalent in human cancers [22,24]. Therefore, understanding the biological role of copines could add to our understanding of how cancer progresses, as well as provide insights into new therapeutic interventions.

## Notes

### Competing Interest Statement

The authors have declared no competing interest.

### Summary of Updates

This version of the manuscript has been revised to update the following: Re-wrote abstract and edited more of the text overall, removed author Bridget K. Plude, removed GFP-CpnD data during amoeba migration, added cAMP stimulation and GFP-CpnD translocation data, added cell size phenotype suppresion after PI3K inhibitor treatment.

### Funding

- National Institutes of Health: R15GM078089-03
- Central Michigan University, Department of Biology
- Central Michigan University, College of Science and Engineering

